# *Pseudomonas aeruginosa* Alkyl Quinolone Response is dampened by Enterococcus faecalis

**DOI:** 10.1101/2024.10.23.619906

**Authors:** Maggie M. Fink, Abigail A. Weaver, Dharmeshkumar Parmar, Jon E. Paczkowski, Lingyun Li, Maggie K. Klaers, Ella A. Junker, Elizabeth A. Jarocki, Jonathan V. Sweedler, Joshua D. Shrout

## Abstract

The bacterium *Pseudomonas aeruginosa* is an opportunistic pathogen that can cause lung, skin, wound, joint, urinary tract, and eye infections. While *P. aeruginosa* is known to exhibit a robust competitive response towards other bacterial species, this bacterium is frequently identified in polymicrobial infections where multiple species survive. For example, in prosthetic joint infections (PJIs), *P. aeruginosa* can be identified along with other pathogenic bacteria including *Staphylococcus aureus, Enterococcus faecalis,* and *Corynebacterium striatum.* Here we have explored the survival and behavior of such microbes and find that *E. faecalis* readily survives culturing with *P. aeruginosa* while other tested species do not. In each of the tested conditions, *E. faecalis* growth remained unchanged by the presence of *P. aeruginosa,* indicating a unique mutualistic interaction between the two species. We find that *E. faecalis* proximity leads *P. aeruginosa* to attenuate competitive behaviors as exemplified by reduced production of *Pseudomonas* quinolone signal (PQS) and pyocyanin. Reduced alkyl quinolones is important to *E. faecalis* as it will grow in supernatant from a quinolone mutant but not P. aeruginosa wildtype in planktonic culture. The reduced pyocyanin production of *P. aeruginosa* is attributable to production of ornithine by *E. faecalis*, which we recapitulate by adding exogenous ornithine to *P. aeruginosa* mono-cultures. Similarly, co-culture with an ornithine-deficient strain of *E. faecalis* leads *P. aeruginosa* to yield near mono-culture amounts of pyocyanin. Here, we directly demonstrate how notorious pathogens such as *P. aeruginosa* might persist in polymicrobial infections under the influence of metabolites produced by other bacterial species.

**Importance:** While we now appreciate that many infections are polymicrobial, we understand little of the specific actions between a given set of microbes to enable combinatorial survival and pathogenesis. The bacteria *Pseudomonas aeruginosa* and *Enterococcus faecalis* are both prevalent pathogens in wound, urinary tract, and bacteremic infections. While *P. aeruginosa* often kills other species in standard laboratory culture conditions, we present here that *E. faecalis* can be reliably co-cultured with *P. aeruginosa.* We specifically detail that ornithine produced by *E. faecalis* reduces the Pseudomonas Quinolone Signal response of *P. aeruginosa*. This reduction of the Pseudomonas Quinolone Signal response aids *E. faecalis* growth.

## Introduction

Many infections are polymicrobial in nature, including those commonly associated with skin wounds, female or male urinary tract, prosthetic joints, and the lung of individuals with cystic fibrosis lung (1–3). Understanding the interspecies interaction between these polymicrobial communities is likely critical to understanding the survival of pathogens in such environments to include mechanisms of antibiotic resistance and evasion of host immune responses. The bacterium *Pseudomonas aeruginosa* is commonly identified in a variety of infections that are polymicrobial (3–8). *P. aeruginosa* virulence factors and toxic attributes are well documented in the literature (9, 10). While *P. aeruginosa* is well-known to exhibit competitive behavior towards other bacterial species (11–15), *P. aeruginosa* does not always dominate. For example, many studies have characterized competitive interactions between *P. aeruginosa* and *Staphylococcus aureus* (16, 17), as both are frequently co-isolated in infections of the lungs and skin and the ratio of *P. aeruginosa* to *S. aureus* does not necessarily change over time (18). Representative interactions between *P. aeruginosa* and other microbes have yet to receive the same research attention. For example, *P. aeruginosa* co-occurrence with *Enterococcus faecalis* (4–6, 19–23) is common as well, but little research has investigated the interaction between *E. faecalis* and *P. aeruginosa*. There are not clear hallmark outcomes associated with *E. faecalis* and *P. aeruginosa* co-infection. *E. faecalis* and *P. aeruginosa* have been specifically identified together in wound infections (4, 20, 24–27), catheter-associated urinary tract infection (CAUTI)(28), periodontal disease (29), and prosthetic joint infection (3, 30). With the rising concern about antibiotic resistance for the Enterococci and Pseudomonads, and high incidence of these pathogens among nosocomial infections, understanding the co-occurrence of these two specific bacterial species is especially important.

Here we examined *P. aeruginosa* in pairwise combination with a select handful of other bacteria that can be present in prosthetic joint infection (PJI). We find that *E. faecalis* survives with *P. aeruginosa* under conditions that outcompete all other tested species. We show that these two species can be co-cultured in planktonic culture and biofilms without compromising the growth of either species. Additionally, we demonstrate an effect of *E. faecalis* on *P. aeruginosa* Pseudomonas quinolone signal production, as well as the synthesis of pyocyanin, a *P. aeruginosa* virulence factor.

These results highlight the ability of *E. faecalis* to dampen the competitive response of *P. aeruginosa*. We specifically identify and confirm a role of the amino acid ornithine in mediating this unique interaction. In addition to potential clinical applications, investigating the ability of *E. faecalis* to interfere with established *P. aeruginosa* alkyl quinolone (AQ) mediated pathways can further our understanding of the complex milieu of factors elicited and sensed by *P. aeruginosa* as part of polymicrobial infections.

## Results

### P. aeruginosa does not exhibit killing towards E. faecalis in co-culture

We were interested to better understand interactions between different bacterial species that have been identified in infection environments like prosthetic joint infection (PJI) (3, 31). We took an approach to characterize responses from a series of pairwise culture experiments that tested *P. aeruginosa* with other PJI microbes. We selected *S. aureus, C. striatum,* and *E. faecalis,* as representative PJI co-isolates. Each species was grown in rich Mueller Hinton broth, alone and in co-culture with *P. aeruginosa.* We found that only *E. faecalis* exhibited measurable CFUs when co-cultured with *P. aeruginosa* (Figure 1A) while *S. aureus* and *C. striatum* had no viable cell count. We were surprised that *E. faecalis* exhibited no significant difference in CFUs from its mono-culture control (Figure 1A). We also tested *E. coli* as a canonical laboratory control and found, as expected, that *E. coli* did not survive in co-culture with *P. aeruginosa.* In contrast, *P. aeruginosa* growth was largely unaffected by the inclusion of any competing species in these experiments as differences in *P. aeruginosa* CFUs from these different co-cultures were minor or insignificant in comparison to monoculture conditions (Figure 1B). While our results are in general agreement with several prior studies reporting that *P. aeruginosa* is generally antagonistic against competing microbes (11, 13, 16, 32, 33), we were struck by the magnitude of difference between the survival of *E. faecalis* in comparison to the killing of *S. aureus, C. striatum,* and *E. coli* measured in our experiments. We were interested to better understand the cooperative relationship between *P. aeruginosa* and *E. faecalis*.

**Figure 1.**
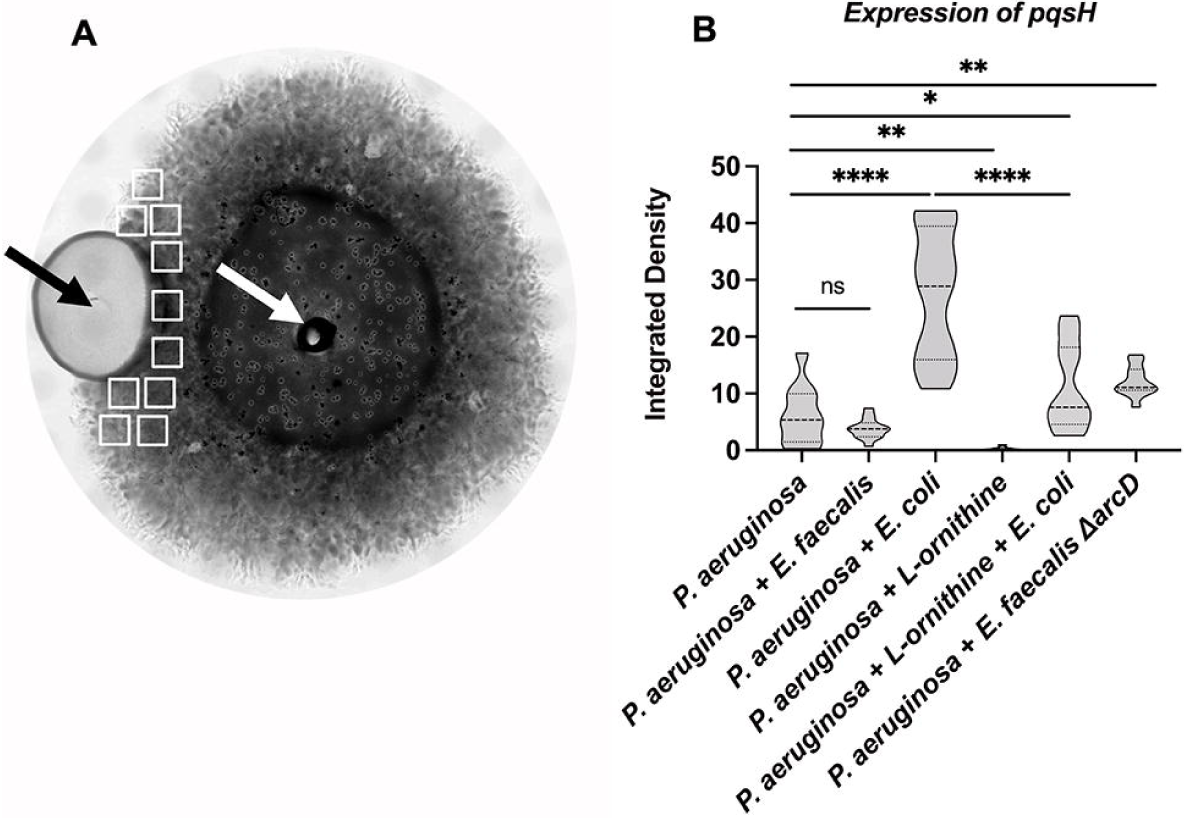
Colony forming units (CFUs) for bacterial species in monoculture and coculture with *P. aeruginosa*, show differences in survival after 24 hours in Mueller Hinton broth. **A**. *S. aureus, C. striatum,* and *E. coli* each have no viable CFUs after being cultured with *P. aeruginosa. E. faecalis* CFUs in coculture with *P. aeruginosa* remain unchanged from monoculture (limit of detection (LOD) = 1). **B.** *P. aeruginosa* CFUs have a slight decrease when cultured with *S. aureus* and *C. striatum* and are unchanged when cocultured with *E. faecalis* and *E. coli.* CFUs were determined from 3 replicates dilutions from the countable plate assay for the range of 10^0^ – 10^-9^ dilution. Pairwise comparisons by Welch’s t-test are indicated on the plot: ns= P > 0.05. *=P ≤ 0.05, ****= P ≤ 0.0001.

### Alkyl quinolone production by P. aeruginosa in colony biofilms is diminished near E. faecalis

One response elicited by other microbes upon *P. aeruginosa* can be measured in the production of heterocyclic aromatic 2-alkyl-4(1*H*)-quinolones (AQs)(13, 34–36). AQs such as 2-heptyl-3-hydroxy-4-quinolone (*Pseudomonas* Quinolone Signal; PQS) and 2-heptyl-4-hydroxyquinoline-N-oxide (HQNO) are known to be produced by *P. aeruginosa* in response to competition with other bacterial species. We have previously shown that *P. aeruginosa* exhibits earlier production of AQs when grown in proximity to other species such as *E. coli* or *S. aureus* (13). We hypothesized that *E. faecalis* would elicit a reduced AQ response, to enable survival in coculture with *P. aeruginosa*.

We examined a series of colony biofilm assay experiments inoculated with *P. aeruginosa* alone or side-by-side with *E. faecalis*. Separate assays included *P. aeruginosa* side-by-side with *E. coli* as an AQ-inducing control (13). To spatially assess relative abundance of *P. aeruginosa* AQ production in response to *E. faecalis* we used matrix-assisted laser desorption/ionization mass spectrometry imaging (MALDI-MSI)(37, 38). A representative spatial heatmap rendering of our results shows greater abundance of PQS near an *E. coli* colony, but not near *E. faecalis*, as compared to *P. aeruginosa* alone (Figure S1). Overall, we assessed PQS as well related quinolones HQNO, HHQ, and their nine carbon side-chain variants C9-PQS, NQNO and NHQ. Intensity profiles for each molecule produced by *P. aeruginosa* near to *E. coli* or *E. faecalis* were assessed relative to amounts produced by *P. aeruginosa* alone (Figure 2). For each of these AQs, we find 2-8× reduced signature is present when *P. aeruginosa* is proximal to *E. faecalis.* As expected, *E. coli* elicited an opposite response as 1.5-2.5× increased signal of PQS, C9-PQS, HQNO, and NQNO was detected above mono-culture levels. We perceived the reduced AQ response near *E. faecalis* as a lack of *P. aeruginosa* antagonism and hypothesized that *E. faecalis* surviving with *P. aeruginosa* is a result of directly altering the AQ response. We directly confirm the importance of reduced AQ production to *E. faecalis* survival as *E. faecalis* will grow when incubated with spent supernatant of a *P. aeruginosa ΔpqsA* AQ deficient strain but not with supernatant of *P. aeruginosa* wildtype (Figure S2).

**Figure 2.**
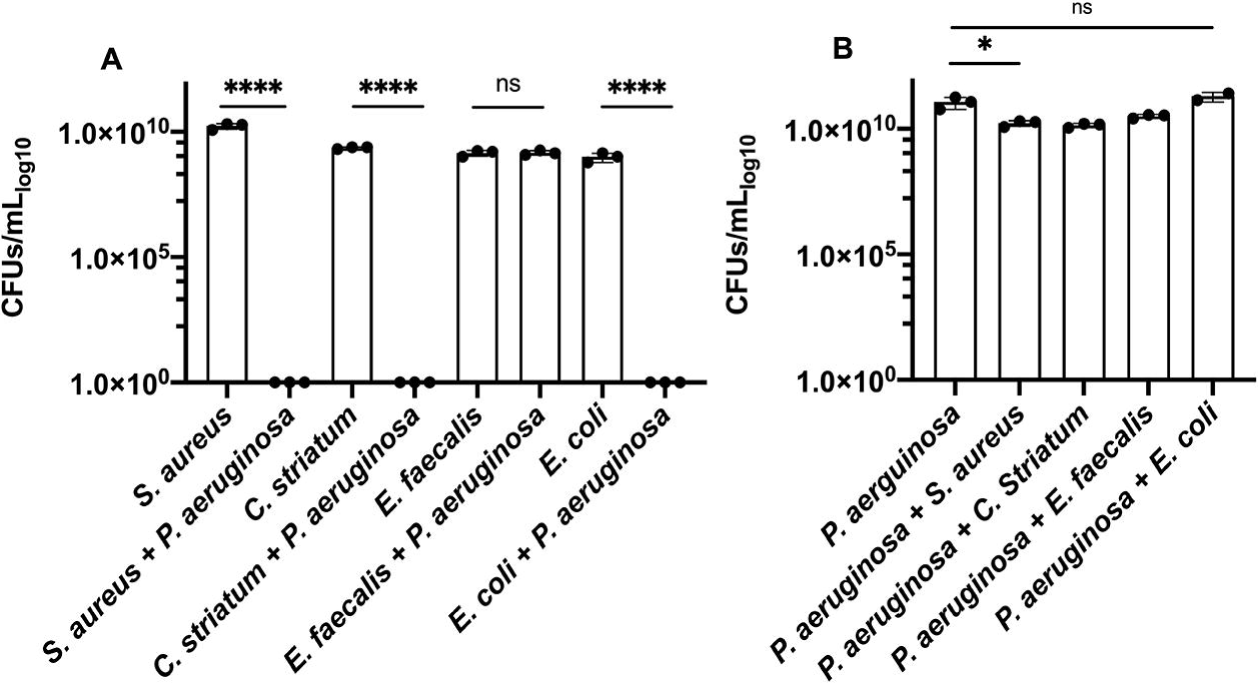
Alkyl quinolone production in *P. aeruginosa* biofilms is reduced by the presence of *E. faecalis.* The amount of alkyl quinolones and pyocyanin detected by MALDI-MSI relative to *P. aeruginosa* alone (i.e. 1.0) indicates *E. coli* and *E. faecalis* have opposite effects on alkyl quinolone production. Error bars show ± one standard deviation. **A.** Presence of *E. coli* elicits an increased PQS, C9-PQS, HQNO and NQNO response while *E. faecalis* reduces production for each. **B.** Additionally, pyocyanin production increased in the presence of *E. coli* (10.3× *P. aeruginosa* alone) but decreased near *E. faecalis* (0.59× *P. aeruginosa* alone).

### E. faecalis reduces expression of the P. aeruginosa alkyl quinolone biosynthetic gene pqsH

We speculated that differences in *P. aeruginosa* AQ abundance observed with *E. faecalis* should result from a substantive shift in *P. aeruginosa* gene expression. The AQ biosynthetic pathway is complex and distinct branch points are known for several well-studied *P. aeruginosa* quinolones (34, 39). We focused our attention on the gene *pqsH,* the final gene required to synthesize the quorum sensing signals PQS and C9-PQS.

We imaged a fluorescent protein transcriptional reporter for *pqsH* and found patterns consistent with our MALDI-MSI results. In 48h biofilm colonies, a basal level of *P. aeruginosa* P*_pqsH-sfgfp_* fluorescence is observed (Figure 3B). Inclusion of *E. faecalis* near *P. aeruginosa* leads to a reduction of P*_pqsH-sfgfp_* expression that is not statistically significant but the spatial variability is reduced; while proximity to *E. coli* increased expression of P*_pqsH-sfgfp_* more than 4.0× (Figure 3B). In these side-by-side colony biofilm assays, any influence of co-culture was spatially dependent as expression of P*_pqsH-sfgfp_* in areas of the *P. aeruginosa* biofilm opposite from *E. faecalis* or *E. coli* were not significantly different (Figure S3). We find equivalent trends in planktonic culture where co-culture with *E. faecalis* leads to reduced P*_pqsH-sfgfp_* expression (67% of mono-culture), while co-culture with *E. coli* increased expression of P*_pqsH-sfgfp_* to 150% of mono-culture (Figure 4A).

**Figure 3.**
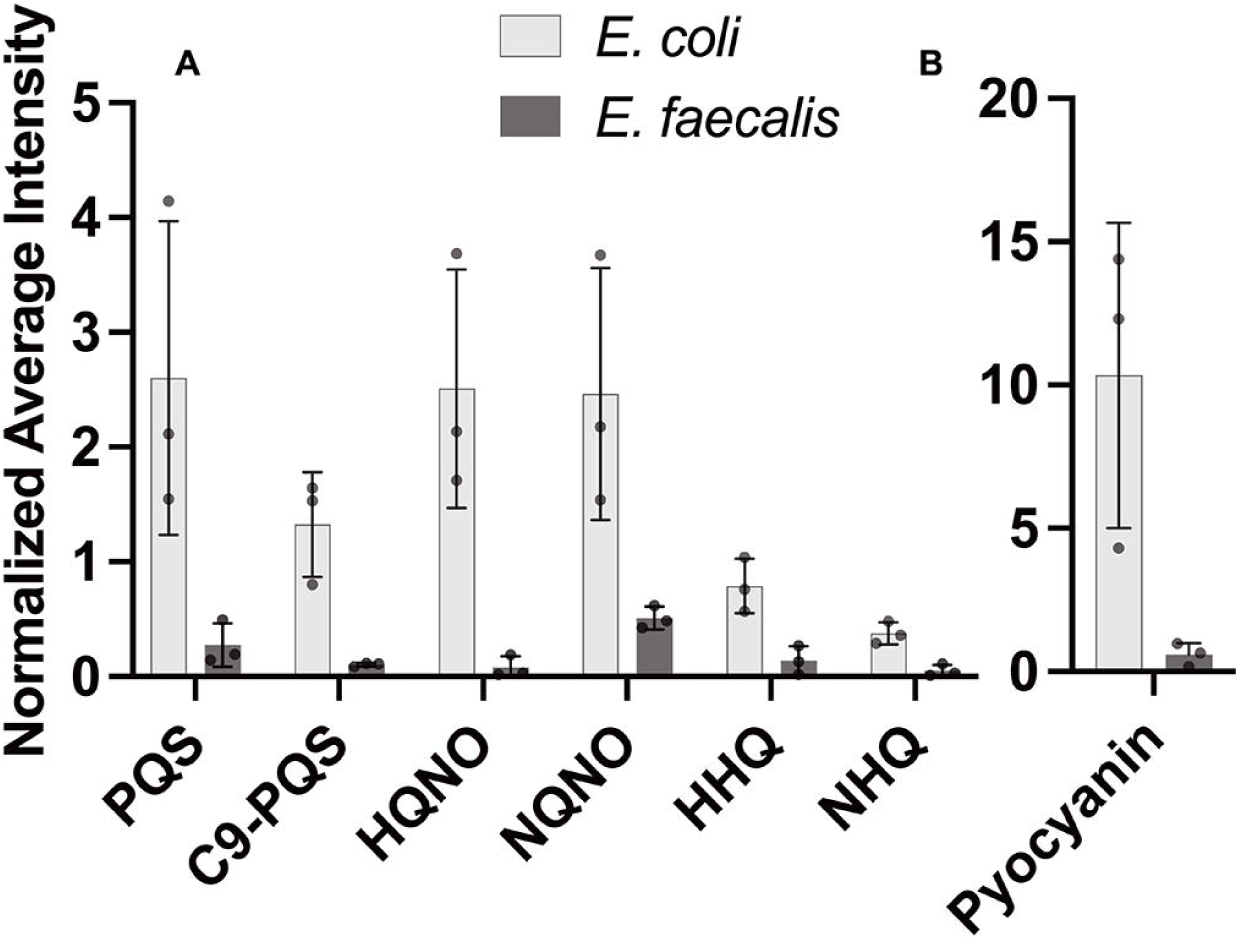
Alkyl quinolone gene expression is reduced in proximity to *E. faecalis*. Fluorescence intensity of P*_pqsH-sfgfp_* was determined after 48 hours incubation. **A.** A representative brightfield image shows the colonies of *E. faecalis* and *P. aeruinogsa* where their sites of inoculation are noted by the black and white arrows, respectively. **B.** No significant difference of *pqsH-sfgfp* was observed between *P. aeruginosa* and *E. faecalis* while *E. coli* significant increased expression. *E. faecalis* Δ*arcD* increased expression of *pqsH-sfgfp*. The addition of 12 mM L-ornithine decreased *pqsH-sfgfp* in monoculture and in coculture with *E. coli.* Integrated densities were acquired from 10 sample areas (white rectangles) on each plate for three biological replicates each for which results were statistically different by one-way ANOVA (P ≤ 0.0001). Pairwise comparisons by Welch’s t-test are indicated on the plot: ns= P > 0.05, *=P ≤ 0.05, **= P ≤ 0.01, ****= P ≤ 0.0001.

**Figure 4.**
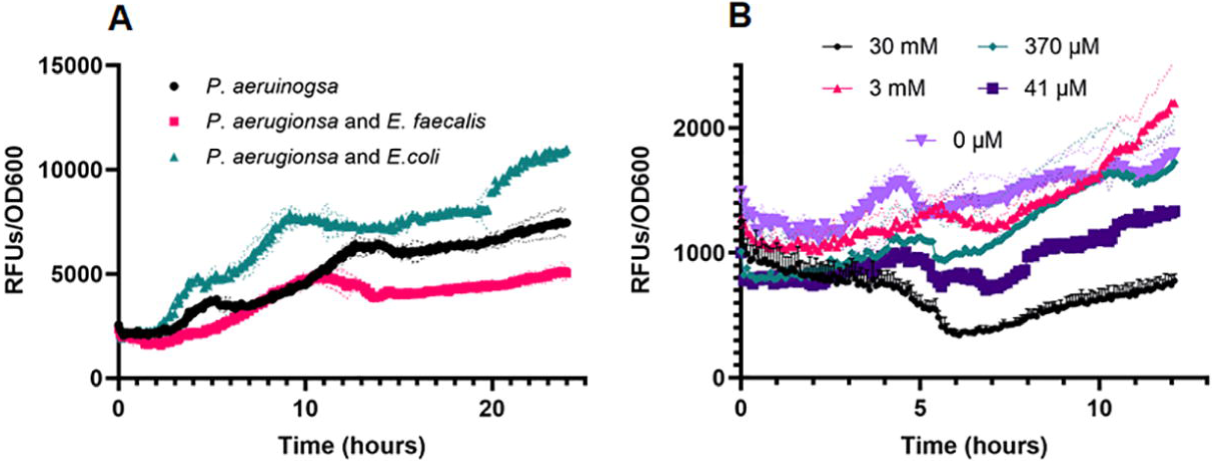
Alkyl quinolone gene expression of *pqsH* is reduced in planktonic co-culture with *E. faecalis* or with exogenous ornithine. A. Fluorescence expression of P_pqsH-sfgfp_ is reduced in planktonic co-culture with *E. faecalis* in comparison to *P. aeruginosa* mono-culture or with *E. coli* (where P_pqsH-sfgfp_ expression is elevated). (B) Fluorescence expression of P_pqsH-sfgfp_ is reduced in planktonic co-culture with increasing amounts of L-ornithine. Values represent averages of ≥3 wells each for which reporter fluorescence was normalized to culture density.

Aside from the PQS regulon, several other competitive effectors and virulence factors produced by *P. aeruginosa* are regulated by the acyl homoserine lactone (AHL) quorum sensing regulons, Las and Rhl (40). Thus, we also tested if the presence of *E. faecalis* directly influenced *P. aeruginosa las* and *rhl* regulon responses by imaging reporters of the three robustly expressed QS genes *hcnA* and *rsaL* (Las) and *rhlA* (Rhl)(41, 42) respectively. The addition of *E. faecalis* near *P. aeruginosa* had minimal effect on the expression of genes regulated by either Las or Rhl as judged by the relative fluorescence of P*_hcnA-gfp_*, P*_rsaL-gfp_*, or P*_rhlA-gfp_* reporters (Figure S4). While a minimal effect on P*_hcnA-gfp_* was observed in our spatial plate assays, we judge this due to heterogeneity of cell density as testing of these same P*_hcnA-gfp_*, P*_rsaL-gfp_*, or P*_rhlA-gfp_* reporters in planktonic culture assays showed no difference in expression with inclusion of *E. faecalis* (Figure S5). Collectively, these results indicate that the influence of *E. faecalis* upon *P. aeruginosa* is specific to PQS and the AQs and not other community-level responses as *E. faecalis* does not affect a change in *P. aeruginosa* Las and Rhl QS gene expression.

### Pyocyanin production is attenuated by the presence of E. faecalis

We noted that in some *P. aeruginosa-E. faecalis* co-culture experiments, *P. aeruginosa* qualitatively displayed less of its characteristic blue-green pigmentation (Figure S6A). The predominant blue *P. aeruginosa* pigment is pyocyanin, for which production is driven by the PQS quorum sensing regulon or interaction of *pqsE* with *rhlR* (9, 34, 43–50). Our MALDI-MSI intensity profile data showed 0.59× amount of pyocyanin near *E. faecalis* while proximity to *E. coli* showed a 10.3× increase of pyocyanin over mono-culture (Figure 2). We probed for a direct effect of *E. faecalis* on pyocyanin production.

To assess boundaries of the expected response, we first tested two separate planktonic culture conditions that yield different relative abundance of *P. aeruginosa* pyocyanin. When *P. aeruginosa* monocultures are grown on glutamate as the sole carbon source, pyocyanin production was maximal, while with glucose, pyocyanin production was minimal. Subsequently, with glutamate as the sole carbon source, *E. faecalis* reduced pyocyanin produced by 42% compared to *P. aeruginosa* monoculture and addition of *E. coli* increased pyocyanin production over monoculture by 115% (Figure 5A). While the amounts of pyocyanin produced with glucose as the sole carbon source are much lower for *P. aeruginosa* monoculture, the addition of *E. coli* increases production to 135% of monoculture. With glucose, pyocyanin production by *P. aeruginosa*-*E. faecalis* co-culture is equivalent to monoculture (Figure 5B), indicating *P. aeruginosa* typical PQS response associated with competitive interactions was attenuated. Thus, with both culture conditions, growing *P. aeruginosa* with *E. faecalis* led to low production, which was opposite of the response quantified with *E. coli*.

**Figure 5.**
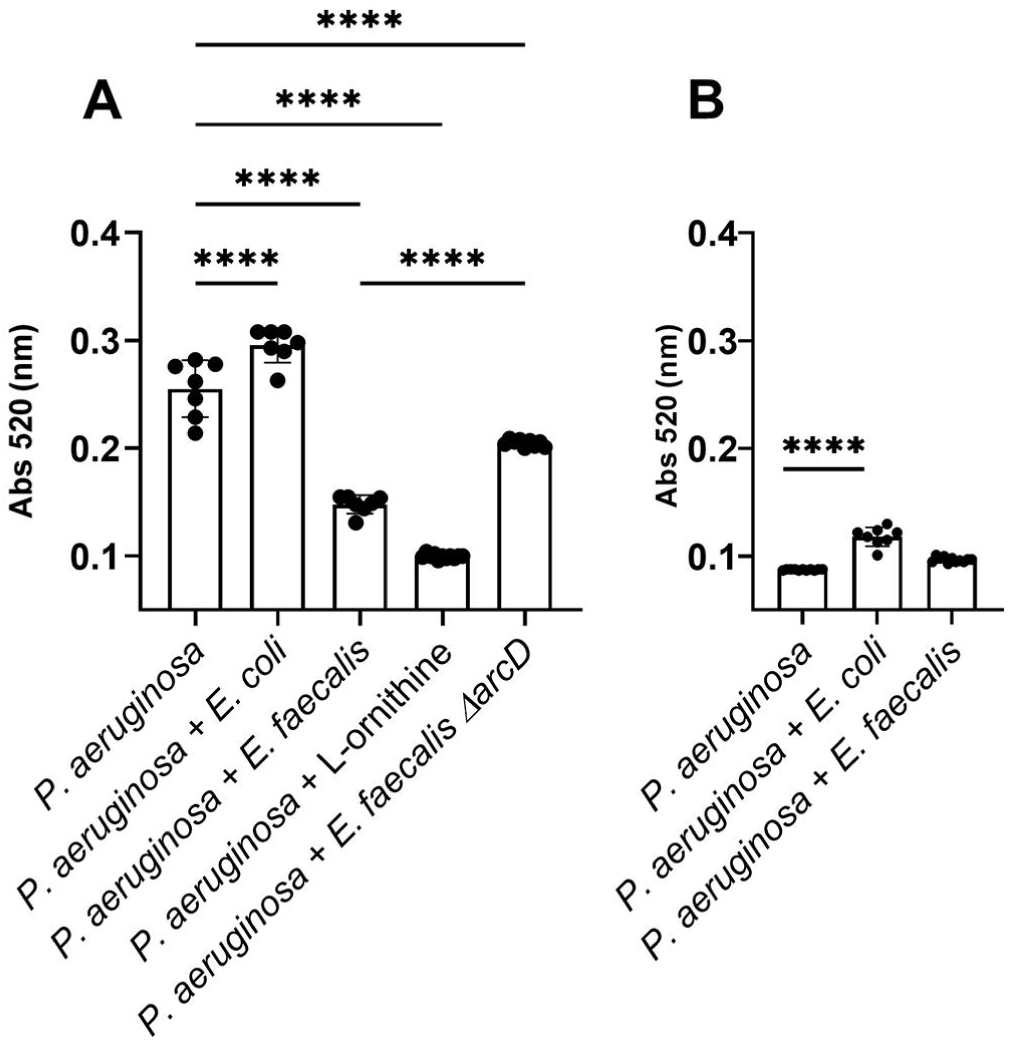
*P. aeruginosa* pyocyanin production is altered by L-ornithine production from *E. faecalis.* Quantification of pyocyanin extracted from *P. aeruginosa* monocultures and cocultures with *E. coli, E. faecalis,* Δ*arcD,* and L-ornithine grown in minimal media containing **(A)** glutamate and **(B)** glucose for which results were statistically different by one-way ANOVA (P ≤ 0.0001). Pairwise comparisons by Welch’s t-test are indicated on the plot: ****= P ≤ 0.0001.

### Ornithine of E. faecalis promotes reduction in PQS response by P. aeruginosa

We were intrigued by a report of Keogh, et al (51) showing that L-ornithine produced by *E. faecalis* altered growth and behavior of *E. coli* in co-culture. We tested if ornithine might be important to the *E. faecalis* and *P. aeruginosa* interactions we observed. First, when equivalent P*_pqsH-sfgfp_* reporter colony biofilm assays to those described above were used, we found that supplementation of growth medium with L-ornithine reduced expression of P*_pqsH-sfgfp_* in *P. aeruginosa* alone to near zero (Figure 3B). Additionally, the addition of L-ornithine led to a reduction in P*_pqsH-sfgfp_* fluorescence even in the presence of *E.coli* (Figure 3B). In planktonic culture, the addition of increasing concentrations of L-ornithine led to greater reduction in P*_pqsH-sfgfp_* fluorescence (Figure 4B).

We further quantified the impact of L-ornithine upon *P. aeruginosa* by confirming a reduction in pyocyanin production. In planktonic culture, we found that increasing amounts of exogenous ornithine (0 - 30 mM) correlated with decreasing levels of pyocyanin (Figure S6B). Using 12 mM of L-ornithine decreased pyocyanin production more than presence of *E. faecalis* (Figure 4B), but concentrations as low as 50 μM also reduced pyocyanin ≥50% (Figure S6B). This effect was specific to L-ornithine, as addition of D-ornithine, or metabolically comparable amino acids L-arginine and L-lysine, did not influence pyocyanin production (Figure S7).

### The ability to produce L-ornithine is key to E. faecalis mediated changes in P. aeruginosa alkyl quinolone pathways

We confirmed that L-ornithine produced by *E. faecalis* is the relevant factor needed to alter *P. aeruginosa pqsH* expression and pyocyanin production testing the *E. faecalis arcD* mutant. *E. faecalis* Δ*arcD* lacks the L-arginine/L-ornithine antiporter and is known to be deficient for ornithine production (51). Unlike wild-type *E. faecalis,* Δ*arcD* elicited increased P*_pqsH-sfgfp_* expression in our colony biofilm model experiments (Figure 3B), thus specifically indicating the importance of ornithine in mediating the interactions between *E. faecalis* and *P. aeruginosa*.

The Δ*arcD* mutant also elicited 1.5x pyocyanin production over wild-type *E. faecalis* (Figure 5A), which was equivalent to 80% of pyocyanin produced by *P. aeruginosa* monoculture. We postulate that L-ornithine may be the dominant factor of multiple made by *E. faecalis* to influence *P. aeruginosa* pyocyanin production.

Lastly, we confirm and quantify production of L-ornithine in planktonic culture using ultra-high performance liquid chromatography (UHPLC) coupled with high-resolution mass spectrometry (HRMS). We find that *E. faecalis* produced L-ornithine in concentrations ranging from 518-539 μM when grown in Mueller Hinton broth (Figure 6A). In coculture with *P. aeruginosa*, L-ornithine was only detected at concentrations ranging from 21 μM - 96 μM (Figure 6A). In *P. aeruginosa* monoculture, L-ornithine was detected in the media at concentrations between 8-13.5 μM. This change in detectable L-ornithine between monocultures and co-culture suggests that *P. aeruginosa* is not just sensing, but also consuming the ornithine produced by *E. faecalis*. While overall ornithine production was substantially lower in minimal medium supplemented with glutamate, we observed similar trends as only *E. faecalis* monocultures show appreciable L-ornithine levels. (Figure 6B).

**Figure 6.**
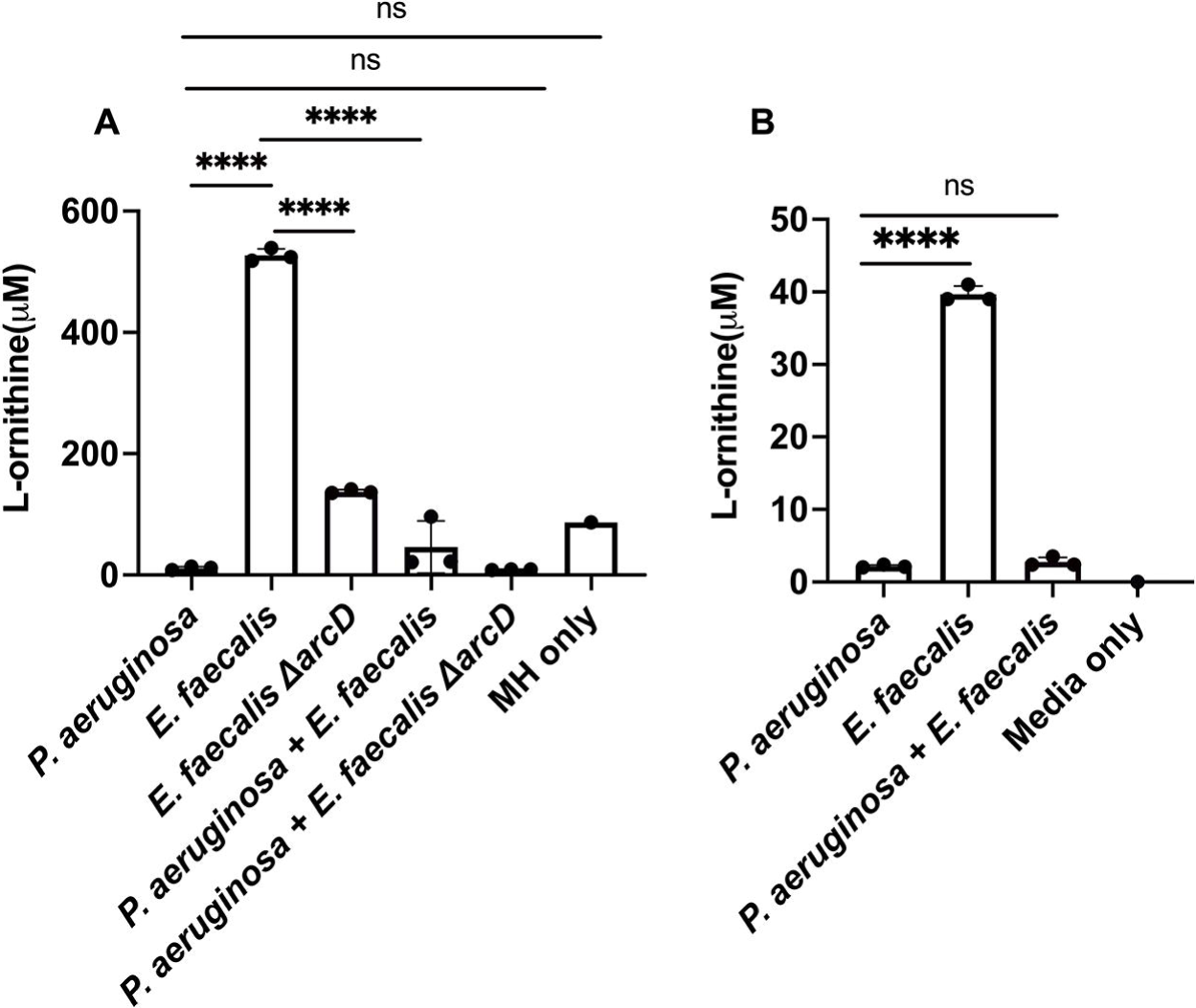
The presence of L-ornithine differs between monocultures of *P. aeruginosa* and *E. faecalis* and co-culture. **A.** In Mueller Hinton broth detectable L-ornithine was decreased in co-culture compared to *E. faecalis* monoculture. *P. aeruginosa* and Δ*arcD* co-culture also had a reduction in L-ornithine compared to Δ*arcD* alone, the amount detected was comparable to L-ornithine already present in the media. **B.** Ornithine was detected in higher abundance from *E. faecalis* monoculture in minimal media supplemented with glutamate compared to *P. aeruginosa* alone and co-culture. Each condition represents three biological replicates for which results were statistically different by one-way ANOVA (P ≤ 0.0001). Pairwise comparisons by Welch’s t-test are indicated on the plot: ns= P > 0.05, ****= P ≤ 0.0001.

## Discussion

Our research sheds light on a unique relationship between *E. faecalis* and *P. aeruginosa.* Specifically, we show a rare example of survival in the presence of *P. aeruginosa* under culture conditions that generally enable *P. aeruginosa* dominance and antagonism. In direct competition, *E. faecalis* dampens *P. aeruginosa* alkyl quinolone responses, which is the opposite of other known examples. Results from this work and prior reports generally show that *P. aeruginosa* alkyl quinolone responses are generally upregulated in the presence of other bacterial species (13, 17). It is not yet clear if the ability of *E. faecalis* to alter *P. aeruginosa* behaviors correlates to competitive or killing responses in infections where both species are present. We do find that absence of *P. aeruginosa* AQs allows for *E. faecalis* growth in planktonic culture. The proximity and range of interaction between *E. faecalis* and *P. aeruginosa* is clearly important as the effect of *E. faecalis* upon alkyl quinolone or pyocyanin production was localized to the area nearest to *E. faecalis*. This finer point was not apparent whatsoever from our planktonic culture results. Nonetheless, this interaction we have characterized between *E. faecalis* and *P. aeruginosa* may provide a useful framework for better understanding specific cross-talk and potential cooperative interactions within polymicrobial communities.

In addition to better understanding bacterial interactions between *E. faecalis* and *P. aeruginosa*, this work also highlights the role of shared metabolites in modulating stress responses and virulence factor production by *P. aeruginosa,* specifically related to the *P. aeruginosa* PQS system. This work demonstrates that an outside signal from *E. faecalis,* L-ornithine, is sufficient to alter the PQS response and reduce the production of the virulence factor, pyocyanin. This is confirmed by showing that an *E. faecalis* Δ*arcD* mutant, which is unable to export L-ornithine, lacks the capacity to limit pyocyanin production by *P. aeruginosa*. Showing this ability of *E. faecalis* to alter the aggressive behavior of *P. aeruginosa* by a specific amino acid represents an important step in understanding *P. aeruginosa* virulence and the potential to develop new treatments for polymicrobial infections.

## Materials and Methods

### Bacterial strains and culture conditions

All strains used in experiments are reported in Table S1. All bacterial strains were routinely grown planktonically for 18 hours in Luria-Bertani (LB) broth at 37°C with shaking at 240 rpm, except as noted for additional specific assays.

Plate assays were prepared using FAB medium supplemented with 12 mM glucose and 1.5% Noble agar. For monoculture plates, *P aeruginosa* was inoculated by pipetting 1 uL of planktonic culture onto the center of the plate. For co-culture biofilms, 1 uL of *E. faecalis* and *E. coli* K-12 were spotted at 12 mm apart from 1 uL *P. aeruginosa*. Plates were inverted and incubated for 24 or 48 hours respectively.

### Viability counts

Colony forming units (CFUs) were determined by using monocultures of each strain inoculated into 1 mL of Mueller-Hinton broth (Sigma-Aldrich) to an OD_600_ of 0.01. Co-culture experiments were inoculated with each individual strain at an OD_600_ of 0.01, for an overall OD_600_ of 0.02 (i.e. at a 1:1 ratio).

Cultures were grown for 24 hours at 37°C shaking at 240 rpm. Triplicate serial dilutions from 10^0^ – 10^-9^ of each monoculture and coculture were done in 1xPBS and 5 uL of each dilution spotted on Brain Heart Infusion agar (Dot Scientific, Burton, MI, USA) agar plates for all species as well as Sabouraud agar (Dot Scientific) for *P. aeruginosa* selection, Mueller-Hinton broth with 2μg/mL Ciproflaxin for *E. faecalis*, Blood agar for *E. coli,* and Mannitol Salt Agar for *S. aureus* and *C. striatum*. Colony forming units (CFUs) were determined for each species in monoculture and coculture using selective media.

### Mass Spectrometry Imaging Analysis

Mass spectrometry images of alkyl quinolones and pyocyanin present on colony biofilm assays were obtained using a FT-ICR mass spectrometer (solariX 7T, Bruker, USA) equipped with matrix-assisted desorption/ionization (MALDI)(52). Additional details are included in the Supplemental Methods.

### Reporter Strain Construction

Standard genetic techniques were used to construct a chromosomal (Tn7) transcriptional reporter strain P*_pqsH-sfgfp_* that fused chromosomal DNA upstream of the *pqsH* gene to sfgfp. Strains, plasmids, and DNA sequences are included in Tables S1, S2 and S3, respectively of the SI Appendix and additional details of construction are included in the Supplemental Methods.

### Fluorescence microscopy of Reporter Strains

Images of *P. aeruginosa* containing P*_pqsH-_*_gfp_, P*_rhlA_*_-gfp_, P*_hcnA_*_-gfp_ and P*_rsaL_*_-gfp_ were obtained using a Leica DM6B upright microscope equipped with a 10× Fluotar objective with simultaneous excitation at 475 nm with emission capture using settings of 525 ± 50 nm. Captured images were 1028 × 916 pixels with a DPI of 144 pixels/in. Grey scale images were obtained using brightfield. Fluorescence intensity was measured using ImageJ (53) where ten equal sized sections of 48×50 pixels each were analyzed individually from three sample replicates for each test condition to determine the resultant integrated density intensity for each condition.

### Pyocyanin Quantification

Pyocyanin quantification extraction was adapted from Frank and Demoss (45).

Additional details included in the Supplemental Methods.

### Ornithine quantification

Ornithine was quantified from planktonic cultures using a hybrid Hydrophilic Interaction Liquid Chromatography (HILIC) coupled with a high-resolution mass spectrometer (HRMS). Additional details included in the Supplemental Methods.

### Data analysis

Plots of data and statistical significance comparison between all tested conditions using one-way ANOVA and pairwise comparisons by Welch’s t-test were generated using GraphPad Prism 10 software.

## Acknowledgments

M.M.F. was funded by NSF Graduate Research Fellowship1841556. J.E.P was funded by National Institute of General Medical Sciences grant R01GM14436101, and J.D.S. and J.V.S. were supported by the National Institute of Allergy and Infectious Diseases grant R01AI113219.

## Author contributions

M.M.F., A.A.W., D.P., J.E.P, J.V.S. and J.D.S. designed research; M.M.F., A.A.W., D.P., J.E.P, L.L., M.K.K, E.A.J., E.A.J. and J.D.S. performed research; J.E.P, J.V.S. and J.D.S. contributed reagents/analytic tools; M.M.F., A.A.W., D.P., J.E.P, and L.L. analyzed data; M.M.F., J.E.P, J.V.S. and J.D.S. funding; and M.M.F., D.P., J.E.P, J.V.S. and J.D.S. wrote and edited the paper.

## Competing interests

The authors declare no competing interest.

## Notes

### Competing Interest Statement

The authors have declared no competing interest.

