## Supplemental Methods, Tables, and Figures for "*Pseudomonas aeruginosa* Alkyl Quinolone Response is dampened by Enterococcus faecalis"

*Mass Spectrometry Imaging Analysis*

Samples for MALDI-MSI analysis were prepared by cutting and removing a center square section of agar from the petri dish, affixing this section to aluminum plates using copper tape, and dehydrating overnight under forced air (1). These dehydrated agar samples were uniformly coated with 2,5-dihydroxy benzoic acid (DHB) (40 mg/mL in 50% Methanol) using M5 overhead matrix sprayer (HTX, USA). The spray parameters were maintained as follows, nozzle height - 40 mm, nozzle temperature - 70 ˚C, lateral nozzle speed - 1000 mm, lateral spacing - 3 mm, matrix flow rate - 0.05 mL/min and nitrogen pressure - 1 bar. Sample surface was coated with DHB in 4 passes with each layer sprayed at 90˚ to the previous one and 10 s drying time between passes. The plates were stored inside the nitrogen desiccator until analysis.

Mass spectrometry imaging was performed on FT-ICR mass spectrometer (solariX 7T, Bruker, USA) equipped with matrix-assisted laser desorption/ionization (MALDI) source. To facilitate MSI, the samples were scanned using a flatbed document scanner at 1200 DPI. Regions of interest for MSI were selected in FlexImaging (version 5.0, build 89). The MS parameters were set using ftmsControl (version 2.3.0, build 59). The data acquisition was carried out in the positive ion mode with laser size set at “ultra-large”, and MS analyzer operated at 1M size (transient data points). Targeted MS/MS (collision energy 30 eV, isolation width ±1.0 Da) was performed to differentiate PQS (*m/z* 260.165>175.062) and its structural isomer HQNO *(m/z* 260.165>159.067), as well as C9-PQS (*m/z* 288.195>175.062) and NQNO (*m/z* 288.195>159.067). Separately, imaging data in a broad mass range (*m/z* 100-1600) was acquired to detect other metabolites, such as pyocyanin. The MSI data was processed in the SCILS lab (Version 2023a Pro). Extracted ion profiles were generated using ±16 ppm extraction window around the observed *m/z*. Extracted ion intensities have been represented using false fire color.

Normalized intensity for analyzed regions were calculated as: {Average intensity from the inoculation zone *second species* }/{Average intensity from the sampling region}.

The normalized average intensity value of 1 (y-axis) indicates the average level in the sampling region. The sampling region is the whole area and includes the intensities from *P. aeruginosa* zone. Normalized average intensity values <1 indicate a relative decrease in the metabolite abundance, and values >1 indicate an increase in the abundance.

*Reporter Strain Construction*

Standard genetic techniques were used to construct chromosomal (Tn7) transcriptional reporter strains to be used as indicators of *P. aeruginosa* AQ or AHL quorum sensing pathway activity. Strains, plasmids, and DNA sequences are included in Tables S1, S2 and S3, respectively. All sequences were designed starting with the PAO1 sequence as a reference strain as annotated on pseudomonas.com (2). A transcriptional reporter for *pqsH* was generated by first amplifying *P. aeruginosa* chromosomal DNA roughly 560 bp upstream of *pqsH* utilizing using the primer pairs listed in Table S3. All primers and gBlocks were obtained from IDT. PCR reactions utilized AllTaq Master Mix (Qiagen) and products cloned with super-folder green fluorescent protein (sfGFP) by isothermal assembly into pUC18-mini-Tn7T-Gm using NEBuilder HiFi Assembly Master Mix and transformed into *E. coli* DH5α (New England Biolabs).

A transcriptional short-half life reporter for *hcnA* was generated by first amplifying *P. aeruginosa* chromosomal DNA previously assessed as regulating transcription of *hcnABC* in *P. fluorescens* (3) using primers in Table S3. A 280 bp fragment was generated by restricting with EcoRI and XbaI, which was cloned into AvrII and EcoRI restricted DNA of pJDS5 to create pJDS116.

Plasmid DNA was transformed by heat-shock into *E. coli* SM10 cells. Conjugational mating was used promote uptake of Tn7 transposons into *P. aeruginosa* (4). Colonies were screened on LB plates supplemented with 100 µg/ml gentamicin or 100 µg/ml tetracycline.

*Fluorescence microscopy of Reporter Strains.*

*P. aeruginosa* harboring chromosomally inserted $P_{pqsH}$-gfp, $P_{rhlA}$-gfp, $P_{hcnA}$-gfp and $P_{rsaL}$-gfp transcriptional fluorescent reporters were imaged to judge the impact of *E. faecalis*, *E. coli*, or exogenous ornithine on relative levels of gene expression. Some of these assays utilized an *E. faecalis* Δ*arcD* mutant (Table S1) that is derived from the *E. faecalis* OG1RF background. (We find the OG1RF wildtype to have equivalent growth to that of EfPJI-A5; c.f. Figure 1 and Figure S8). Images were obtained using a Leica DM6B upright microscope equipped with a 10× Fluotar objective with simultaneous excitation at 475 nm with emission capture using settings of 525 ± 50 nm. Grey scale images were obtained using brightfield. From each region of interest, ten replicate sections (48 × 50 pixels) were sampled to determine the integrated fluorescence density as measured using ImageJ (5).

Quantification of reporter expression for planktonic cultures over time using a microplate reader (Biotek, Synergy H1). *P. aeruginosa* wildtype harboring $P_{pqsH}$-gfp, $P_{rhlA}$-gfp, $P_{hcnA}$-gfp or $P_{rsaL}$-gfp reporter constructs was added to 200 µL FAB-glucose in black 96-well clear bottom plates (Agilent) to an initial OD of 0.01. Absorbance at 600 nm and fluorescence at excitation of 488 nm with emission capture at 525 nm were measured every 10 minutes for 24 hours at 37°C. The plate was shaken for 10 seconds before each read.

*E. faecalis monoculture with P. aeruginosa supernatant growth conditions*

The *P. aeruginosa* wildtype and Δ*pqsA* strains were each grown planktonically in 6mL FAB medium supplemented with 12 mM glucose while *E. faecalis* EfPJI-A5 was grown planktonically in Luria-Bertani (LB) broth. These planktonic cultures were incubated for 20 hours at 37°C with shaking at 240 rpm. Spent-culture supernatant was harvested from the *P. aeruginosa* cultures was by filter-sterilizing the resultant supernantant after 10 minutes centrifugation.

Using clear 96-well plates, 100 µL of *P. aeruginosa* wildtype or Δ*pqsA* spent culture supernatant was added to 100 µL sterile FAB-glucose. Non-supernatant controls contained 100 µL sterile FAB. Wells were then inoculated with 5 µL *E. faecalis* that was diluted 1 in 5 from the pre-grown planktonic culture. The plate was grown at 37°C with 10 seconds shaking every 10 minutes prior to each optical density measurement at 600 nm using a microplate reader (Biotek, Synergy H1).

*Pyocyanin Quantification*

Pyocyanin quantification was adapted from Frank and Demoss (6). To extract pyocyanin from planktonic cultures, 6 mL planktonic cultures were grown for 24 hours, then centrifuged at 12,000 rpm for 20 minutes at 4°C. Supernatants were collected, filtered with 0.22μm sterile filters, and mixed with 3 mL of chloroform. After vortexing for 20 seconds and allowing the aqueous and organic phases to separate, the top aqueous layer was removed. 20% total volume of 0.1N HCL was added to the chloroform fraction containing the pyocyanin and allowed to settle for 10 minutes. The aqueous solution containing crude pyocyanin extract was removed and absorbance was measured at 520 nm in a black 96-well glass bottom plate (Aligent) using a microplate reader (Biotek, Synergy H1).

*Ornithine quantification*

Monocultures and cocultures were grown as described for determining CFUs. At 24h cultures were spun down at 10,000 rpm for 20 minutes and supernatant was filtered using 0.22 μm PES syringe filters (Avantor). A hybrid Hydrophilic Interaction Liquid Chromatography (HILIC) coupled with a high-resolution mass spectrometer (HRMS) method was used for ornithine quantitative analysis. A high-throughput and direct sample extraction method was developed for cell-free supernatant sample analysis. Briefly, 100 μL of filtered supernatant samples were directly extracted using 900 μL acetonitrile/water mixture (9:1) and vortexed for 15 s. The precipitated proteins were separated from the extraction solvent using centrifugation (2 min, 10,000 × g). The supernatant was directly used for HILIC-HRMS analysis. For HILIC-HRMS high-throughput analysis, the instrumental system included a Vanquish liquid chromatography system coupled with a high-resolution QE Orbitrap mass spectrometer (ThermoFisher) operating in the positive electrospray-ionization (ESI) mode with a heated ion source (HESI) running at parallel reaction monitoring (RPM) mode. Analyte separation was achieved using a HILIC column (Raptor Polar X, 2.7 µm, 100 mm x 2.1 mm, Restek) as the stationary phase and solvent A and B as mobile phases. For gradient elution, mobile phase A was 0.5% formic acid, 1 mM ammonium formate in water, and mobile phase B was 0.5% formic acid, 1 mM ammonium formate in acetonitrile: water (90:10). The flow rate was 0.40mL/min, and the column was maintained at 35 °C. The binary gradient was initially at 96% mobile phase B and decreased to 15% mobile phase B over 4.5 min. After a hold at 5% mobile phase B from 4.6-5.4 min, the mobile phase composition was returned to 96% mobile phase B in 0.1min and maintained for another 2 min to equilibrate the column. The sample injection volume was 2 µL. An external calibration curve was used for quantifying the ornithine in the samples with quality control samples to ensure the system stability was qualified for each batch. We used a standard spike-recovery rate in this analysis. For quantification, a 5 parts per million (ppm) accurate mass window was used for peak integration of the high-resolution MS/MS data acquired using the PRM method. The limit of quantification of 0.30 µM ornithine was achieved using this method. Data were acquired and processed with TraceFinder 5.0 software (ThermoFisher).

Table S1: Strains used in this study

| Strain | Select Characteristics | Reference |
|---|---|---|
| *Escherichia coli* | | |
| DH5α | High-efficiency for cloning | New England Biolabs |
| SM10 | Conjugational competent donor; λpir | (7) |
| K12 | Standard wild-type strain | (8) |
| *Pseudomonas aeruginosa* | | |
| PAO1C | Wildtype Laboratory Strain, ATCC Collection Strain 15692 | (9-12) |
| Δ*pqsA* | PAO1C Δ*pqsA* | (13) |
| wt-P*rhlA*-gfp | 15692 $P_{rhlA}$-*gfp* reporter; $Tc^r$ | (14) |
| wt-P*rsaL*-gfp | 15692 $P_{rsaL}$-*gfp*(AAV) reporter; $Tc^r$ | (15) |
| wt-P*hcnA*-gfp | 15692 $P_{hcnA}$-*gfp*(AAV) reporter; $Tc^r$ | This study |
| wt-P*pqsH*-gfp | 15692 $P_{pqsA}$-gfp reporter; $Gm^r$ | This study |
| *Enterococcus faecalis* | | |
| EfPJI-A5 | Clinical isolate from prosthetic joint infection | (16) |
| 10103 | OG1RF Δ*arcD* | (17) |
| *Staphylococcus aureus* | | |
| SaPJI-C | Clinical isolate from prosthetic joint infection | (16) |
| *Clostridium striatum* | | |
| CsPJI-A24 | Clinical isolate from prosthetic joint infection | (16) |

Table S2: Plasmids used in this study

| Plasmid | Select Characteristics | Reference |
|---|---|---|
| pUC18-mini-Tn7T-Gm | $Ap^r$; $Gm^r$ on mini Tn7T | (18, 19) |
| pTNS2 | $Ap^r$; R6K replicon; TnsABC+D transposition pathway | (18, 19) |
| pJDS5 | p-miniCTX-$P_{rsaL}$::*gfp*(ASV); $Tc^r$ | (15) |
| pJDS116 | p-miniCTX-$P_{hcnA}$::*gfp*(ASV); $Tc^r$ | This study |
| pJDS151 | pUC18-mini-Tn7T-Gm-$P_{pqsH}$-sfgfp | This study |

Table S3: Primers and DNA used in this study

| Primer | Sequence |
|---|---|
| PpqsH_EcoRI_UP-F | GATCCCCCGGGCTGCAGGAATTCTTCAGCACGATCCACTCGTAG |
| PpqsH_sfGFP_UP-R | TCTTCTCCTTTGCTCATCCGTTGCTCCTTAGCAGCGGCATC |
| EcoRI-PhcnA-F | GAATTCCGTCGCTGTCGGTGAACGAA |
| XbaI-PhcnA-R | TCTAGATTGCCCTTTCATCCGTGAGA |
| PpqsH-sfGFP_DN-F | GCTAAGGAGCAACGGATGAGCAAAGGAGAAGAACTTTT |
| sfGFP_HindIII_DN-R | CGCGAGGTACCGGGCCCAAGCTTTTACGCTGCAAGGGCGTAATTTTC |
| sfgfp-F | ATGAGCAAAGGAGAAGAACTTTT |
| sfgfp-R | TACGCTGCAAGGGCGTAATTTTCG |
| sfgfp gblock | ATGAGCAAAGGAGAAGAACTTTTCACTGGAGTTGTCCCAATTCTT<br>GTTGAATTAGATGGTGATGTTAATGGGCACAAATTTTCTGTCCGT<br>GGAGAGGGTGAAGGTGATGCTACAAACGGAAAACTCACCCTTAA<br>ATTTATTTGCACTACTGGAAAACTACCTGTTCCGTGGCCAACACT<br>TGTCACTACTCTGACCTATGGTGTTCAATGCTTTTCCCGTTATCC<br>GGATCACATGAAACGGCATGACTTTTTCAAGAGTGCCATGCCCG<br>AAGGTTATGTACAGGAACGCACTATATCTTTCAAAGATGACGGG<br>ACCTACAAGACGCGTGCTGAAGTCAAGTTTGAAGGTGATACCCT<br>TGTTAATCGTATCGAGTTAAAGGGTATTGATTTTAAAGAAGATGG<br>AAACATTCTTGGACACAAACTCGAGTACAACTTTAACTCACACAA<br>TGTATACATCACGGCAGACAAACAAAAGAATGGAATCAAAGCTAA<br>CTTCAAAATTCGCCACAACGTTGAAGATGGTTCCGTTCAACTAGC<br>AGACCATTATCAACAAAATACTCCAATTGGCGATGGCCCTGTCCT<br>TTTACCAGACAACCATTACCTGTCGACACAATCTGTCCTTTCGAA<br>AGATCCCAACGAAAAGCGTGACCACATGGTCCTTCTTGAGTTTG<br>TAACTGCTGCTGGGATTACACATGGCATGGATGAGCTCTACAAA<br>GCAGCGAACGACGAAAATTACGCCCTTGCAGCGTAA |

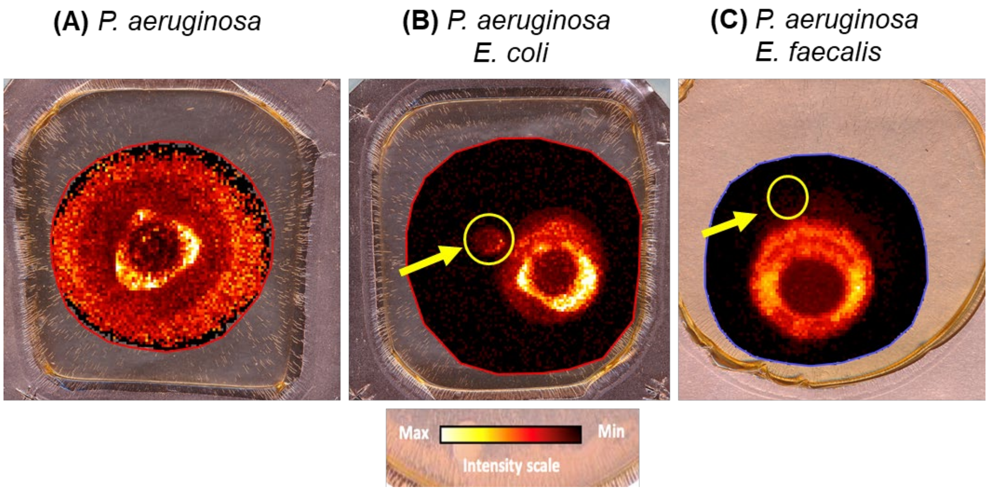


**Figure S1.** Spatial MSI intensity profiles for PQS show distinct localization patterns for 48-hour colony biofilms of **(A)** *P aeruginosa* alone, near **(B)** *E. coli,* and near **(C)** *E. faecalis.* Images show the PQS intensity heatmap overlayed on the camera image of the agar assay. Each assay shows spatial variability of *P. aeruginosa* produced PQS throughout the colony biofilm. The highest levels appear as a ring surrounding the center of inoculation. When *P. aeruginosa* is co-cultured adjacent to *E. coli* (yellow circle), PQS is detected into the region where *E. coli* has grown. When *P. aeruginosa* is co-cultured adjacent to *E. faecalis* (yellow circle), no apparent increase in PQS is observed in this region.

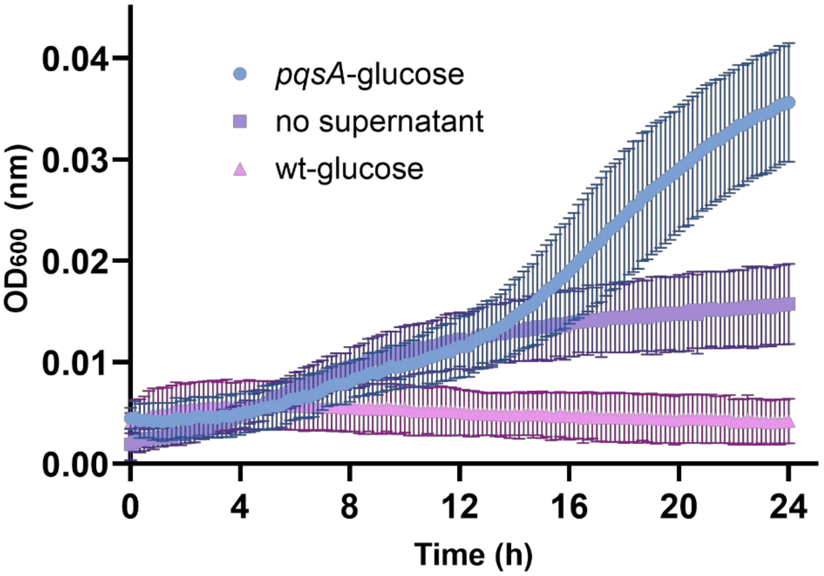


**Figure S2.** Absence of the *P. aeruginosa* PQS synthesis cascade (Δ*pqsA*) enables *E. faecalis* monoculture growth. Addition of filtered supernatant from spent *P. aeruginosa* Δ*pqsA* enables while *E. faecalis* growth while growth is inhibited with spent *P. aeruginosa* wildtype supernantant. A control condition with unamended medium allows marginal growth of *E. faecalis.* Error bars show standard deviation from ≥8 biological replicates.

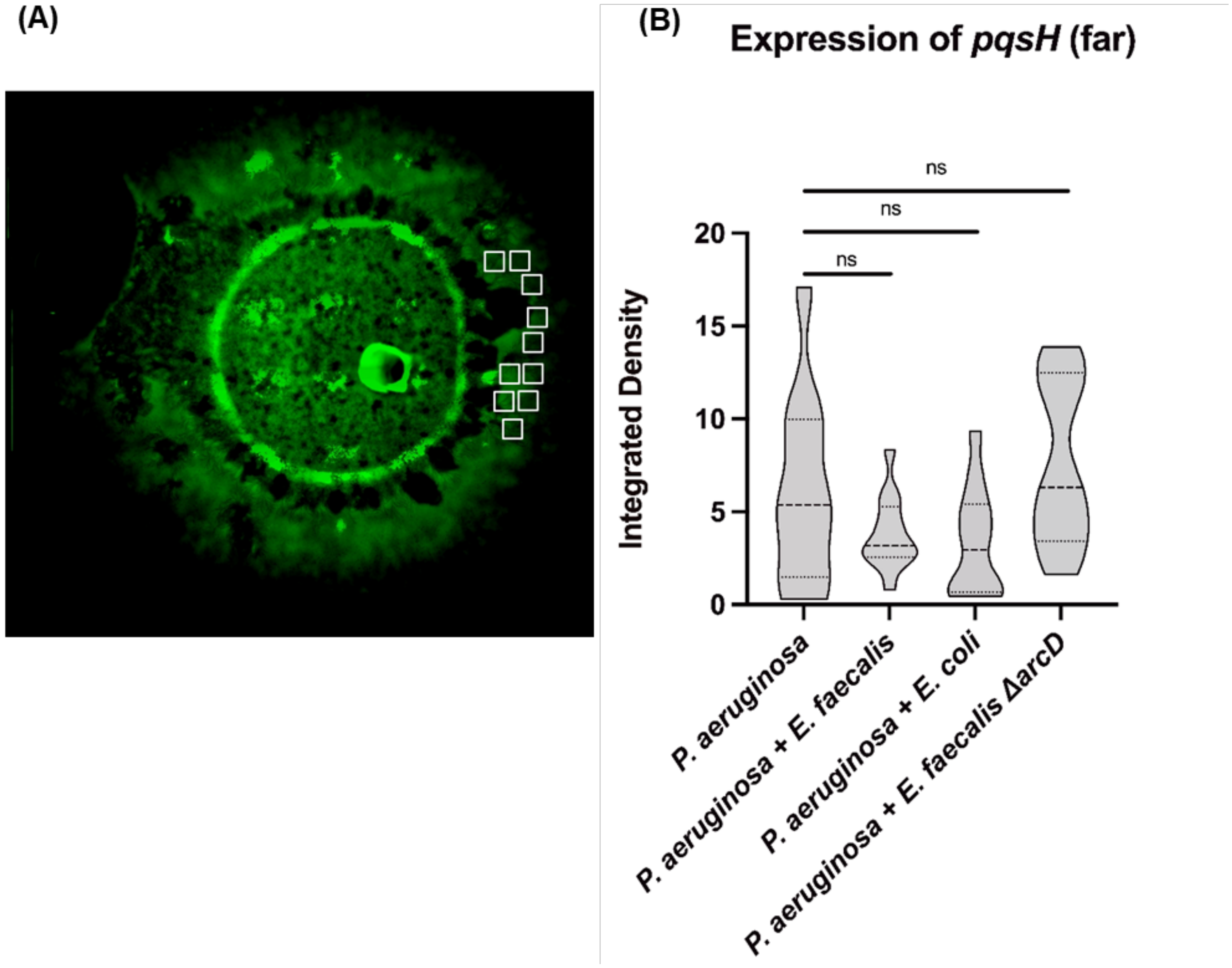


**Figure S3.** Expression of $P_{pqsH\text{-}sfgfp}$ was measured in *P. aeruginosa* biofilm edge opposite the area of contact with *E. coli* or *E. faecalis.* (This is directly comparable with results shown in Figure 3 that show expression directly adjacent to *E. faecalis*.) **A.** The representative assay shows green fluorescence of $P_{pqsH\text{-}sfgfp}$ by *P. aeruginosa* when *E. faecalis* is adjacent on the left with the “opposite region”. **B.** Integrated densities were acquired from 10 sample areas (white rectangles) on each plate for three biological replicates each for which results were statistically different by one-way ANOVA ($P \leq 0.0001$). Pairwise comparisons by Welch’s t-test are indicated on the plot: ns= $P > 0.05$.

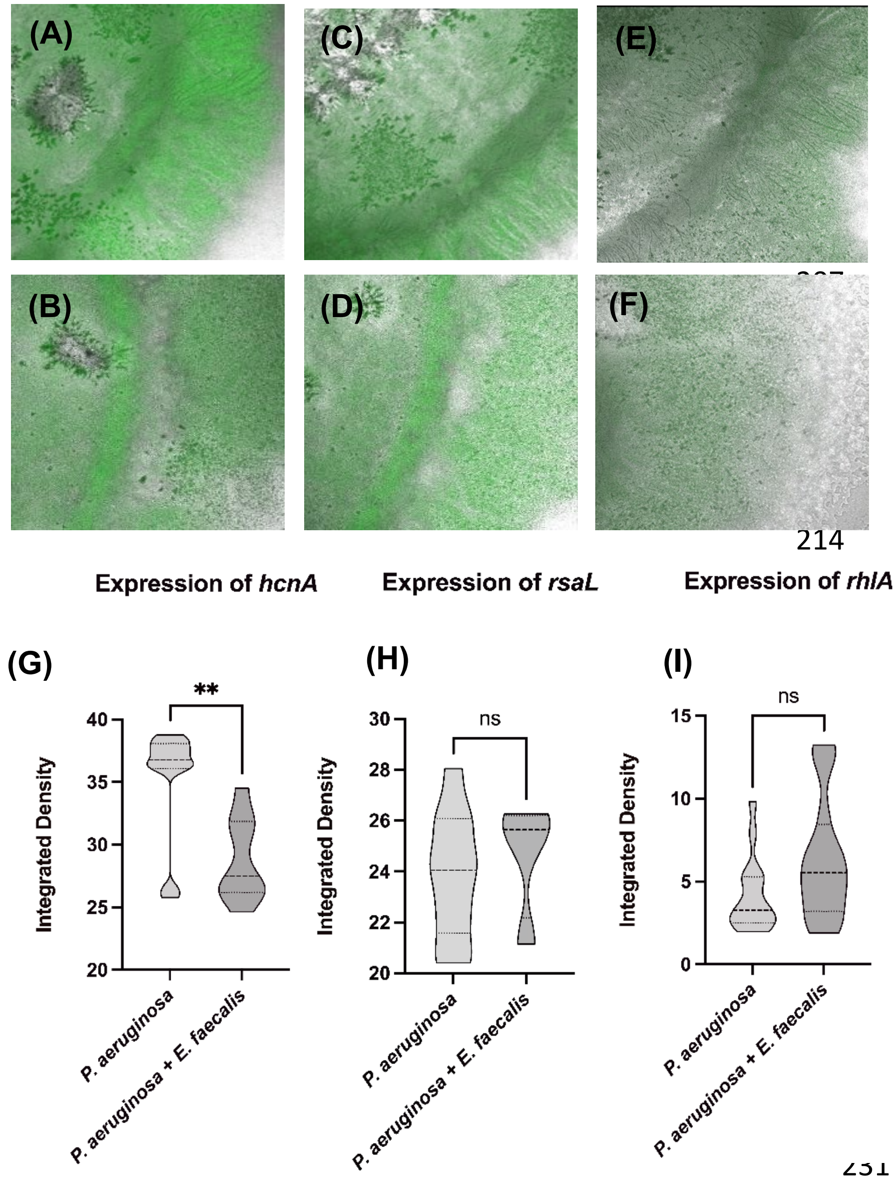


**Figure S4. A-F.** AHL quorum sensing gene expression is unaffected in proximity to *E. faecalis.* Fluorescence expression of $P_{hcnA\text{-}gfp}$ by *P. aeruginosa* near colony edge is equivalent in surface growth assays for **(A)** *P.a*. monoculture or **(B)** near *E. faecalis*. Expression of $P_{rsaL\text{-}gfp}$ by **(C)** P.a. monoculture or near **(D)** *E. faecalis.* Expression of $P_{rhlA\text{-}gfp}$ by **(E)** *P.a*. monoculture or **(F)** near *E. faecalis*. **G-I.** Quantification of fluorescence corresponding to expression of respective genes. Each reporter strain condition was replicated on three separate plates and comparisons by Welch's t-test are indicated on the plot: ns= $P > 0.05$, **= $P \leq 0.01$.

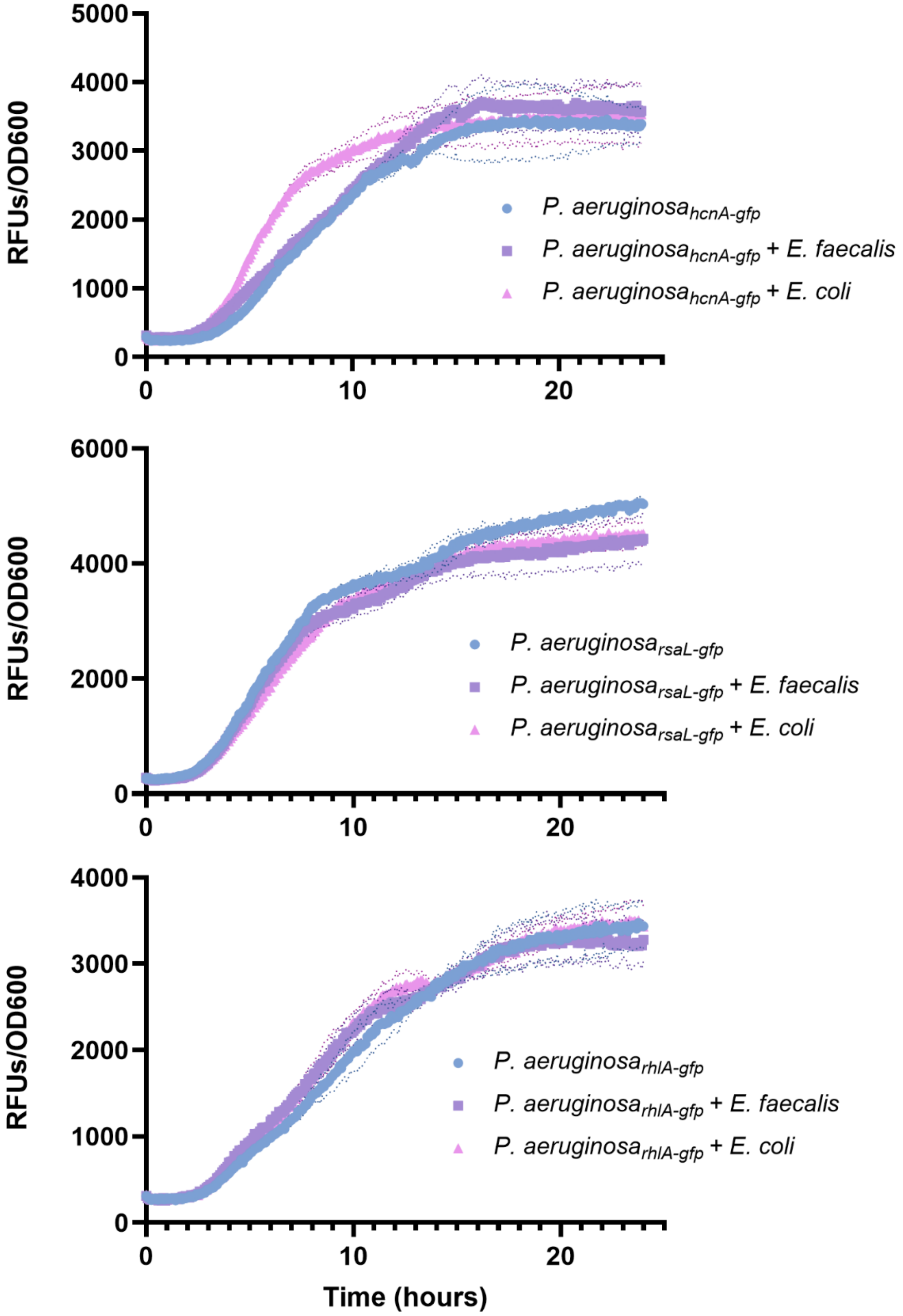


**Figure S5.** AHL quorum sensing gene expression is unaffected in planktonic co-culture with *E. faecalis.* Fluorescence expression of (A) $P_{hcnA\text{-}gfp}$ (B) $P_{rsaL\text{-}gfp}$ (C) $P_{rhlA\text{-}gfp}$ by *P. aeruginosa* is normalized to culture density over time.

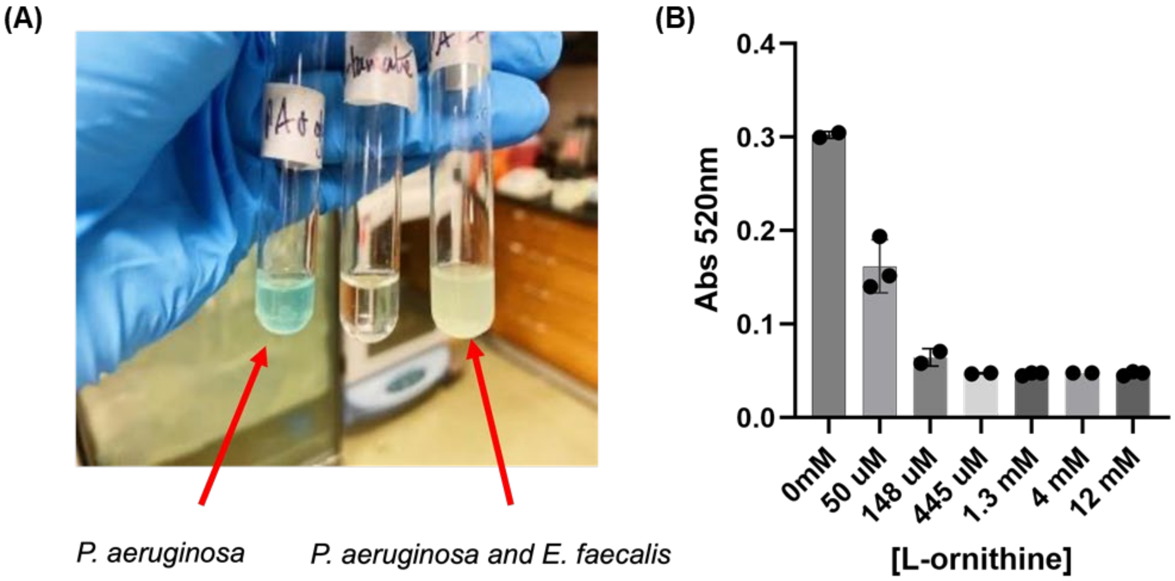


**Figure S6. A.** The characteristic blue pigment of pyocyanin is present in *P. aeruginosa* monoculture but is absent in co-culture with *E. faecalis.* **B.** Exogenous addition of increasing concentrations of L-ornithine to *P. aeruginosa* cultures decreases pyocyanin production (by Frank and Demoss method (6)). Error bars show ± one standard deviation.

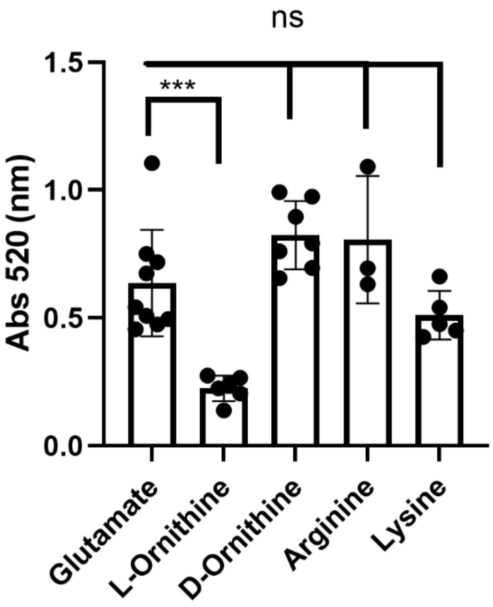


**Figure S7.** Pyocyanin production is not changed by D-ornithine or other metabolically and structurally related amino acids arginine and lysine. Error bars show standard deviation of ≥3 biological replicates for which results were statistically different by one-way ANOVA ($P \leq 0.0001$). Pairwise comparisons by Welch's t-test are indicated on the plot: ns= $P > 0.05$, ***= $P \leq 0.001$.

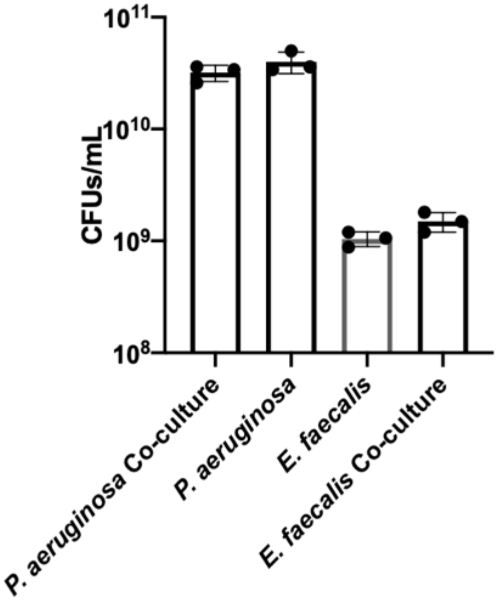


**Figure S8.** Colony forming units (CFUs) for *P. aeruginosa* and *E. faecalis* OG1RF from monoculture and coculture experiments after 24 hours in Mueller Hinton broth. CFUs were determined from 3 replicates dilutions. Error bars show ± one standard deviation.
